# A Series of Composited Tumor DNA Reference Materials Containing Three Genes and Ten Mutation Positions for CNV and SNV Detection

**DOI:** 10.1101/2023.05.04.538185

**Authors:** Weijian Fan, Runyu Shi, Hongyun Zhang, Cuicui Li, Jiawen Zhang, Shuixiu Su, Ping Wu, Meifang Tang

**Affiliations:** Clinical laboratory of BGI Health, BGI-Shenzhen, Shenzhen 518083, China

**Keywords:** Reference materials, droplet digital PCR, sequencing panels, SNV and CNV

## Abstract

Processes in clinic for tumors diagnosis and treatment need reference materials (RMs) to evaluate and calibrate. However, no RMs can provides properties of copy number variation (CNV) and single nucleotide variants (SNV) of genes *EGFR, HER2, MET, PIK3CA, KRAS, BRAF, NRAS* simultaneously. In this study, we used commercial cell lines to construct a series of tumor RMs containing property mentioned above. Furthermore, we evaluated their stability, homogeneity, and commutability by droplet digital PCR and next generation sequencing technology. The results showed that, for tumor CNV gDNA RM, the copy number is 7.3 copies/μL (*EGFR*), 5.3 copies/μL (*HER2*) and 8.2 copies/μL (*MET*). For tumor 5% SNV gDNA RM, the mutation frequency of each mutation position showed as follow: *EGFR*-E746A750 (24.6%), *EGFR*-L858R (5.8%), *EGFR*-T790M (5.5%), *EGFR*-G719S (6.6%), *PIK3CA*-E545K (4.7%), *PIK3CA*-H1047R (5.8%), *KRAS*-G13D (8.2%), *KRAS*-G12D (6.5%), *BRAF*-V600E (4.6%), *NRAS*-Q61K (8.5%). All variable coefficient (CV) of tumor gDNA RM for homogeneity were less than 7%, that of CNV+SNV ctDNA RM were less than 17%. Besides, the CV for commutability of the all types of RMs were less than 17%. These RMs can be applied into a wide range type of sequencing panels and provides a closer simple background.

## Introduction

Cancer is the most important leading cause of death until 2021(Siegel et al. 2021). In 2018, there were 18 million new cancer diagnosed cancer cases(Mattiuzzi and Lippi 2019) and it was predicted that there will be 32.1 million of diagnosis cases in 2050(Pilleron et al. 2021). Among various types of tumor, Lung & bronchus cancer, breast cancer and colon & rectum cancer was the top three tumor types causes death in 2021(Siegel et al. 2021). They share several mutations in genes such as *EGFR, MET* and *HER2* [*BREE2*] (Du and Lovly 2018). Currently, these genes as well as *PIK3CA, KRAS, BRAF* and *NRAS* are taken as biomarkers to facilitate clinic diagnosis of cancer(Nicholson et al. 2001; Matsumoto et al. 2017; Lee and Loree 2019; Moore et al. 2020). These biomarker genes have been broadly applied into cancer therapy and prevention.

World Health Organization (WHO) reported that 50% of cancer cases could be prevented by effective and inexpensive technologies such as initial diagnosis, screening. Thus, an increasing attention is paid into the detection technologies of tumor including diagnosis and early screening. With the growing requirement of tumor screening, regular methodology plays a pivot role.

Nowadays, a range of methods to measure the expression of tumor biomarkers are available and wide-spread. Results from quantitative PCR (qPCR) or digital PCR (dPCR) can provide accurate results of copy number (Wang et al. 2014) and mutation frequency of genes. Researchers or doctors can measure the copy number of cancer biomarkers like *EGFR* from patients’ samples by qPCR and dPCR(Nicholson et al. 2001; Gridelli et al. 2007) and can monitor whether there is the early state of tumor or not. Furthermore, next-generation sequencing (NGS) also contributes the high-throughput sequencing detection of gene mutations(Lee and Loree 2019). making it widely used in clinical diagnosis or cancer-related research. Although these methods play an important role in detections of tumor biomarkers, the quantification of copy number variation (CNV) and the detection of detecting single nucleotide variation (SNV) of genes are still easily being affected by the performance of platforms of detecting platforms or the bioinformatic approaches(Nicholson et al. 2001; Wang et al. 2014; Siegel et al. 2021). Also, the outcomes of detecting instruments may be unreliable owing to their instability. It is necessary to use CNV and SNV reference materials (RMs) to evaluate the performance of different methods(Fay 1967; Hardwick et al. 2017).

DNA RMs are characterized as materials or substance in which one or more property values are sufficiently homogeneous and well established. They can be used for the evaluation and calibration of a measurement method or equipment, or assigning values to materials(Pratt et al. 2009; He et al. 2021).. The pivot function of RMs is to provide a uniform source of stable material that can be used to calibrate different detecting platform, to manage quality control of the detection processes and evaluate the performance of different methods. Thus, RMs help calibrate different measuring systems so that every test proceed is stable. No matter on different detecting instruments or with different technicians, reliable results can be exported(He et al. 2016; Mattiuzzi and Lippi 2019; Siegel et al. 2021). There are currently numerous ways to develop RMs, but the already commercially available RMs have shortages to some extent. The National Institute of Standards and Technology (NIST) has developed SRM2373 and RM8366 for the measurement of *HER2* (*ERBB2*) copy number, *EGFR* and *MET* copy number respectively. However, both the two NIST RMs were merely designed to monitor copy number of genes but mutation rates(He et al. 2019). On the other hand, part of RMs in market are developed in the form of plasmid or PCR product. Although they are convenient to develop and easy to access great mutation rate, the large copy number of gene inside cause strong diffusivity and high proportion of false positive. Some other RMs are developed from clinical patients’ samples, which have advantages since they provide real human genetic background. But clinical patient samples are rare to access and are not easy to produce RMs in mass-scale for their poor homogeneity. Synthetic DNA is a supplement method to develop RMs which is competitive by its quick accessing and ease to get rare mutations. This forms of RMs, however, are various, which poses challenge to quantification. RMs that are composited by clinical patient cells line with interested gene mutation can avoid the shortages of other forms of RMs. Since these RMs’ cell lines are also from clinical sample, they have identical human genetic background to other clinical samples. These RMs can be pre-treated by the same processes of clinical sample, making the quality control of whole processes possible.

In this research, we constructed a series of new RMs by using cell lines from clinical samples. This series of RMs have two forms: genome DNA (gDNA) and circle tumor DNA (ctDNA). They were evaluated from the aspects of homogeneity, stability and commutability by droplet digital PCR (ddPCR) assays and next generation sequencing (NGS) assay. Besides, this series of RMs also contain ten important tumor progression-relative mutation positions, which are *EGFR*-E746_A750, *EGFR*-L858R, *EGFR*-T790M, *PIK3CA*-E545K, *PIK3CA*-H1047R, *KRAS*-G13D, *BRAF*-V600E, *KRAS*-G12D, *EGFR*-G719S, *NRAS*-Q61K. These RMs with good property can be applied into calibration of of ddPCR, quantitative PCR and sequencing panel to quantify tumor biomarkers for cancer detection and diagnosis.

## Materials and Methods

### Cell lines and cell culture

We prepared eleven human cancer cell lines: NCI-H1650, NCI-H1975, NCI-H1944, NCI-H460, NCI-H2087, HCC1954, COLO205, GP2D, HCC827, EBC1 and SW48 to develop tumor SNV and/or CNV gDNA RMs. Those cells lines contained different mutation positions respectively (Table 1) and all of these cell lines contain various copy number information of *EGFR, HER2* and *MET*. These cell lines were bought American Type Culture Collection (ATCC, Manassas, VA) as frozen stocks. These stocks were thawed resume and cultured in our laboratory with standard cell culture protocol.

**Table 1.**
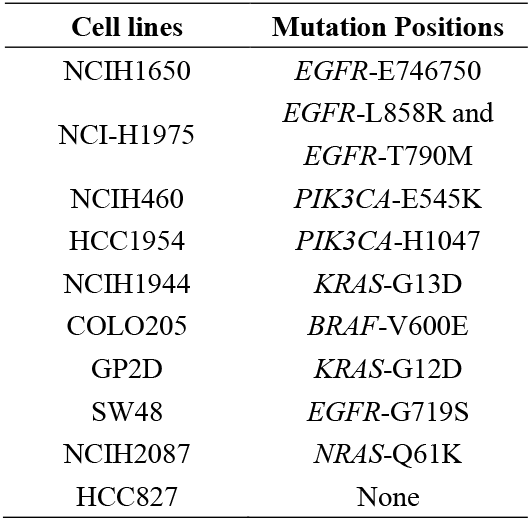
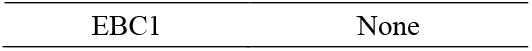
List of Cell lines with information of mutation position and genes in each cell lines

### RMs Preparation

#### DNA extraction

The cells were sub-cultured for 4 to 5 passages, and harvested for DNA extraction when they reached around 90% confluence in ten T-175 cell culture flasks. Genomic DNA samples were extracted using MagPure Buffy Coat DNA Midi KF Kit (MAGEN Cat#D3537-02). All extracted genomic DNA samples were diluted to around 50 ng/μL by TE-4 buffer (10 mmol/L Tris, 0.1 mmol/L EDTA, pH 8.0), and quantified by Qubit 3.0 (Thermo Fisher Scientific-CN). We subsequently used Eppendorf® BioPhotometer to determining the DNA purity of samples based on the absorbance ratio of 260nm and 280nm (A260/A208≈1.8). The molecular mass and integrity of the DNA were assessed by 1% agarose gel electrophoresis.

#### STR Identification

The gDNA extractd from different cell line was authenticated by the human DNA typing kit (platinum) from BGI, Ltd. Co. on a 3730xl DNA Analyzer with a 50-cm capillary array (Thermo Fisher Scientific). Data from the 3730xl was analyzed using GeneMapper ID-X software (version 1.3; Life Technologies). Genome DNA samples were analyzed independently when the cell lines were thawed from frozen stocks and when the cells were sub-cultured for DNA extraction. The short tandem repeat markers will be compared to the values documented by ATCC, to ensure correct cell lines were used.

#### Droplet digital PCR detection

We employed ddPCR assays to quantify copy number of genes and to detect mutation rates of ten mutation position. This experiment was applied on BIO-RAD QX200 droplet digital PCR system and with TaqMan™ fluorescent probe method, following the instruction of BIO-RAD ddPCR™ Copy Number Variation Assays and the ddPCR Applications Guide. The ddPCR reaction systems and procedures referred to Online Resouse 1.

Three and six sets of primers/probes were designed for each target gene and reference genes respectively, to measure the copy number of *EGFR, MET* and *HER2* in cells.

For each single nucleotide variant positions, one primers/probes set was used. The SNV positions we focused on are *EGFR*-E746_A750, *EGFR*-L858R, *EGFR*-T790M, *PIK3CA*-E545K, *PIK3CA*-H1047R, *KRAS*-G13D, *BRAF*-V600E, *KRAS*-G12D, *EGFR*-G719S, *NRAS*-Q61K. All these sets of primer and probe were obtained from Invitrogen, Sangon Biotech (shanghai, China) and BIO-RAD. The details of these sets of primer and probe for CNV and SNV qualification were shown in Online Resouse 2.

#### Cell lines assembling

RMs were prepared according to the copy number of the genes - *EGFR, HER2* and *MET* – and the mutation frequency of the ten mutation positions in the eleven cell lines. We developed five types of tumor DNA RMs. They were tumor 5 % SNV gDNA RM including ten mutation position at mutation rate around 5% (except EGFR-E746_A750), tumor CNV gDNA RM with three genes with amplified gene copy number of genes *EGFR, HER2* and *MET*, and tumor SNV + CNV gDNA RMs with quantitated mutation frequencies of the mutation position of tumor genes (mutation frequency are between 1%-16%) and amplified gene copy number of gene *EGFR, HER2* and *MET*.

The tumor ctDNA RM were prepared from the fragmentated gDNA RM by ultrasonication SN 004924 Covaris. The mutation frequency and copy number of SNV and CNV or tumor ctDNA RM was evaluated by ddPCR. The mutation frequency of the ten mutation positions in 5% tumor SNV ctDNA RM was close to 5%, the properties of SNV of tumor SNV gDNA RM and tumor SNV + CNV ctDNA RM are similar with that of tumor SNV + CNV gDNA RM. More details were shown in Online Resouse 3.

In this research, we chose tumor SNV gDNA RM, tumor CNV gDNA RM and tumor SNV + CNV ctDNA RM as represent to evaluate their performance for the reason that ctDNA are more sensitive to storage environment, which is more representative to the stability of RM.

#### Quantification of CNV and SNV of RMs

RMs’ copy number of the three genes and the mutation frequency of the ten mutation positions were determined by ddPCR in three replicates. The variable coefficient of copy number of each gene was calculated.

### RMs quality control and performance evaluation

#### Concentration and integrity of RMs’ DNA

The DNA concentration of tumor gDNA RMs were determined by two people (P1 and P2) using Qubit™ dsDNA HS assay kit in three biological replicates and two technical replicates.

The integrity of the gDNA in RMs were assessed by agarose gel electrophoresis. 60 ng of gDNA and 250 ng of Lambda DNA-mono cut mix (New England Lab, cat# N3019S) were loaded in different wells of 1% agarose gel in 1x TAE buffer. The gel was run with 220V for 20min. The DNA bands were imaged with Ethidium Bromide staining and were recorded as a digital photograph under UV light. The length of ctDNA fragment was analyzed by agilent 2100.

#### Homogeneity of RMs

To test the homogeneity of RMs, we chose the 5% SNV gDNA RM, CNV gDNA RMs and SNV + CNV ctDNA RM to evaluate the RMs’ intra-batch stability and inter-batch stability. We used ddPCR to detect and compared the mutation frequency of the ten mutation positions and/or copy number of RMs in two biological replicates (Intra-batch) and three technical replicates(batch-to-batch).

#### NGS assay to test the RMs performance in different panels

The tumor ctDNA RM was sequenced in three cancer panels on BGI-Sequencing platform: drug panel, pan-cancer panel, lung cancer_T panel. Plus, 1x Whole-Genome Sequencing (1x WGS, sequenced by DNBSEQ-T7) were adopted to evaluate the Commutability of RMs. The 1x WGS was conducted on BGI-Seq platform.

#### Monitoring of the stability of RMs

RMs were stored in dark place at 4°C for one month and three months respectively, and the copy number of the three genes and the mutation frequency for the ten mutation positions were then detected by ddPCR. The stability across time was described by variable coefficients.

## Results

### Characteristic of tumor cell lines’ genome DNA

Copy number of *EGFR, HER2* and *MET* and mutation frequency of the ten mutation positions in eleven cell lines had been calculated before developing tumor gDNA RMs, the copy number of *EGFR, HER2* and *MET* and mutation frequency of the ten mutation positions in eleven tumor cell lines were quantified. The results were shown in Table 2 and Table 3. According to the genes’ copy number and the mutation frequency of the eleven cell lines, we mixed cells by calculating proportions to prepare tumor CNV gDNA RMs, tumor 5% SNV gDNA RMs and tumor SNV + CNV gDNA RMs.

**Table 2.** Mutation frequency of CNV and SNV. a. Copy number of *EGFR, HER2* and *MET* in the 11 cell lines;

|  | *EGFR* | *HER2* | *MET* |
| --- | --- | --- | --- |
| **NCIH1650** | 12.37 | 1.91 | 2.11 |
| **NCIH1975** | 3.76 | 2.43 | 2.91 |
| **NCIH460** | 0.82 | 2.44 | 2.33 |
| **HCC1954** | 2.36 | 116.3 | 2.4 |
| **NCIH1944** | 2.07 | 2.07 | 1.98 |
| **COLO205** | 1.97 | 2.76 | 1.95 |
| **GP2D** | 1.95 | 2.07 | 1.97 |
| **SW48** | 3.09 | 2.10 | 3.05 |
| **NCIH2087** | 2.80 | 2.41 | 1.90 |
| **HCC827** | 55.4 | 3.59 | 2.9 |
| **EBC1** | 2.07 | 2.06 | 14.5 |

**Table 3.** The mutation frequency (%) of the ten mutation positions in the eleven cell lines.

|  | **EGFR-**<br>**E746_A750** | **EGFR-**<br>**L858R** | **EGFR-**<br>**T790M** | **PIK3CA-**<br>**E545K** | **PIK3CA-**<br>**H1047R** | **KRAS-**<br>**G13D** | **BRAF-**<br>**V600E** | **KRAS-**<br>**G12D** | **EGFR-**<br>**G719S** | **NRAS-**<br>**Q61K** |
| --- | --- | --- | --- | --- | --- | --- | --- | --- | --- | --- |
| **NCIH1650** | 91.6 | 0 | 0 | 0 | 0 | 0 | 0 | 0 | 0 | 0 |
| **NCIH1975** | 0 | 77.3 | 77.7 | 0 | 0 | 0 | 0 | 0 | 0 | 0 |
| **NCIH460** | 0 | 0 | 0 | 50.7 | 0 | 0 | 0 | 0 | 0 | 0 |
| **HCC1954** | 0 | 0 | 0 | 0 | 80.3 | 0 | 0 | 0 | 0 | 0 |
| **NCIH1944** | 0 | 0 | 0 | 0 | 0 | 32.8 | 0 | 0 | 0 | 0 |
| **COLO205** | 0 | 0 | 0 | 0 | 0 | 0 | 67.1 | 0 | 0 | 0 |
| **GP2D** | 0 | 0 | 0 | 0 | 0 | 0 | 0 | 49.8 | 0 | 0 |
| **SW48** | 0 | 0 | 0 | 0 | 0 | 0 | 0 | 0 | 34.4 | 0 |
| **NCIH2087** | 0 | 0 | 0 | 0 | 0 | 0 | 0 | 0 | 0 | 35.4 |
| **HCC827** | 0 | 0 | 0 | 0 | 0 | 0 | 0 | 0 | 0 | 0 |
| **EBC1** | 0 | 0 | 0 | 0 | 0 | 0 | 0 | 0 | 0 | 0 |

### Copy number and mutation frequency of RMs

The copy number of three genes and the mutation rate of ten position of RMs were quantified. The copy number of the three genes were calculated by the ratio of concentration of the target gene and the reference gene.

The results were shown in Figure 1. For the tumor CNV gDNA RM, the copy number of *EGFR* was 7.3 copies/mL with CV of 1.5%, that of *HER2* was 5.3 (CV = 1.7) and *MET* was 8.2 copies/mL (CV = 0.8%). For tumor SNV 5% gDNA RM, the mutation rates of each positions were stated as follow: *EGFR*-E746A750 (the mutation frequency was 24.6%, CV=2.4%, same below), *EGFR*-L858R (5.8%; CV=8.1%), *EGFR*-T790M (5.5%;CV=7.1%), *EGFR*-G719S (6.6%;CV=7.9%), *PIK3CA*-E545K (4.7%;CV=10.1%), *PIK3CA*-H1047R (5.8%;CV=11.2%), *KRAS*-G13D (8.2%;CV=3.9%), *KRAS*-G12D (6.5%;CV=6.4%), and *BRAF*-V600E (4.6%;CV=11.3%), *NRAS*-Q61K (8.5%;CV=5.5%).

**Figure 1.**
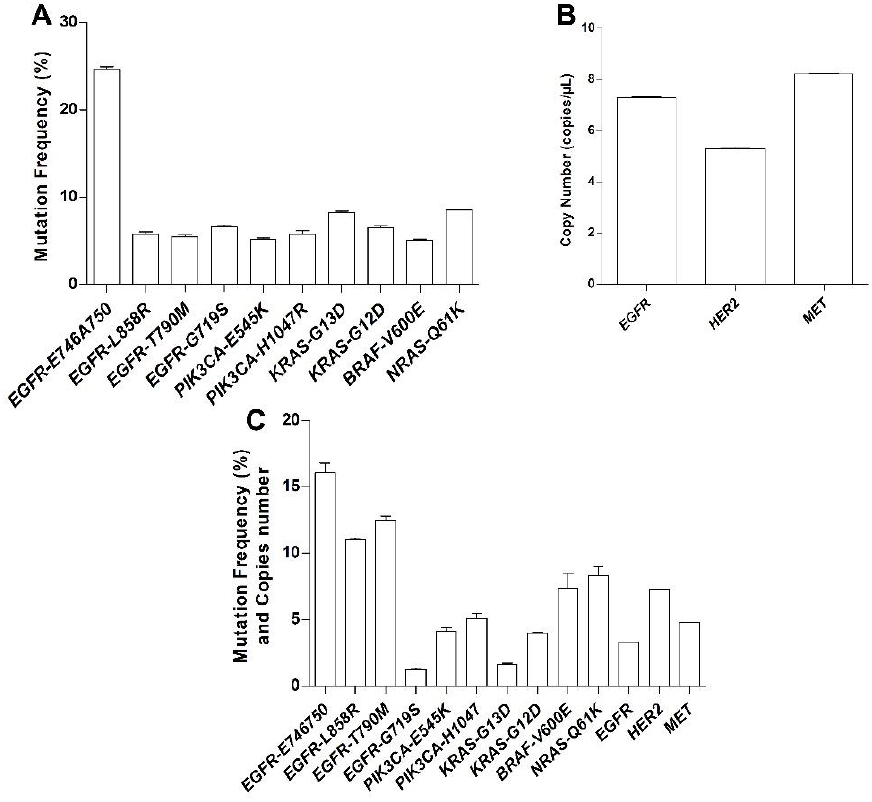
The property of tumor gDNA RMs. a. The mutation frequency of the ten mutation positions of tumor 5% SNV gDNA RMs; b. The copy number of three genes of tumor CNV gDNA RMs; c. The mutation frequency and copy number information of tumor SNV and CNV RMs.

### DNA concentration, Genome DNA integrity and two ctDNA RMs fragment size distribution

According to the copy number of the genes and the mutation frequency of ten mutation positions in the eleven cell lines, we developed two tumor gDNA RMs. The DNA concentration of tumor SNV 5% gDNA RM and CNV gDNA RM were shown in Figure 2. The average concentration of the tumor SNV 5% gDNA RM was 52.4 ng/μL (CV = 6.13%), and the concentration of the tumor CNV gDNA RM was 52.3 ng/μL (CV = 2.36) There was no significant difference between the result from the two operators (P=0.7).

**Figure 2.**
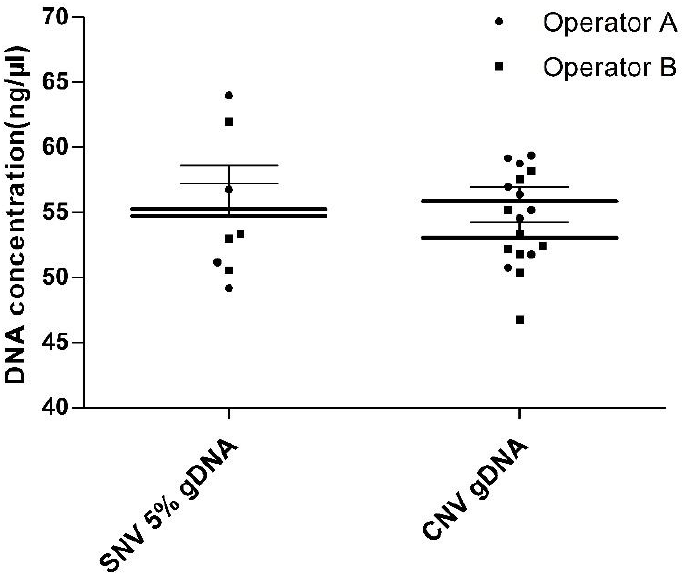
The consistence of DNA concentration of the RMs. For the tumor SNV 5% gDNA RM, two operators quantified the DNA concentration by seven replicates by ddPCR. For the tumor CNV gDNA, two operators conducted ddPCR to quantified the DNA concentration for 17 replicates.

The DNA integrity of tumor SNV 5% gDNA RM and CNV gDNA RM were shown in Figure 3a. Additionally, the DNA fragment size of ctDNA RMs mainly distributed between the range of 159bp to 180bp (Figure 3b), and over 60% fragments were in 100-250bp (Table 4).

**Table 4.**
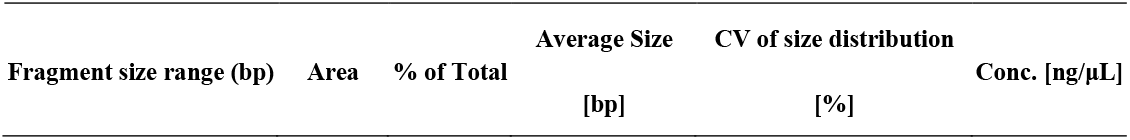
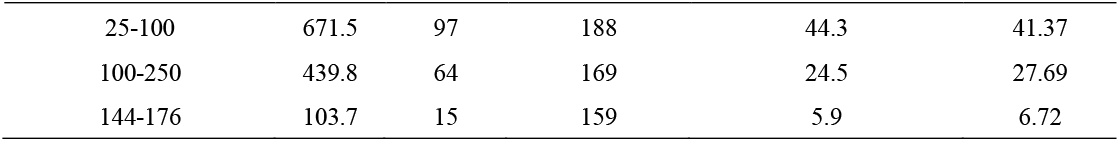
ctDNA fragment distribution ration of tumor SNV RMs

**Figure 3.**
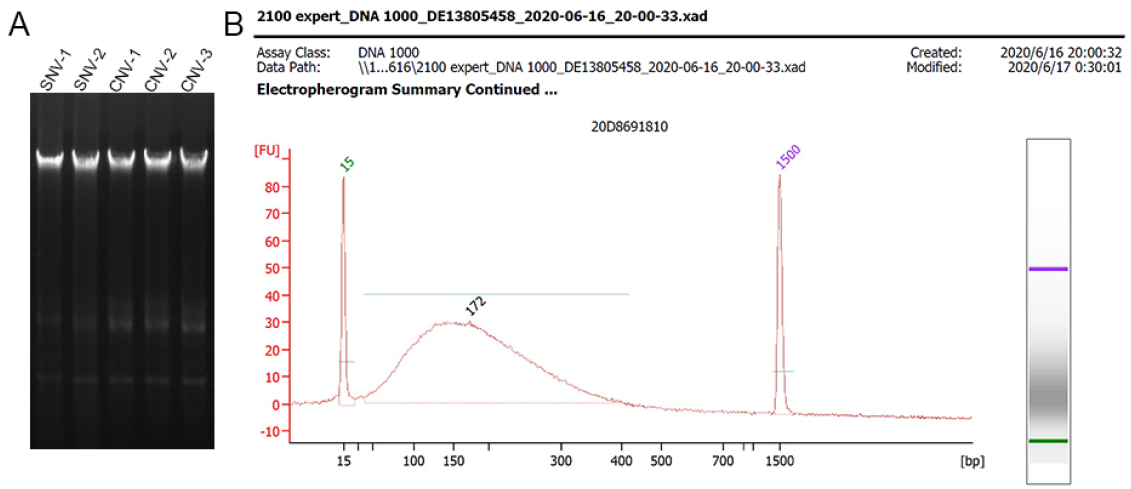
The integrity and DNA concentration of the RMs. The gel electrophoresis image of tumor gDNA RMs (CNV and 5% SNV RMs) in 1% agarose gel electrophoresis, all three lanes were repeats of one gDNA sample; b the DNA fragment size of the tumor SNV + CNV ctDNA RMs.

### Homogeneity

To evaluate the homogeneity of the series of RMs, we selected tumor ctDNA RMs from two different batches and the samples from the same batch to conduct ddPCR. The results were shown in Figure 4. Mutation frequencies of the ten mutation positions were quantitated, their average CVs were then calculated. Intra-batch group shown 6.56% average CV, and that of inter-batch group was 6.48%, indicating a good homogeneity of the tumor SNV 5% RM. The average CV of the tumor CNV gDNA inter- and intro-batches are less than 1.6%, giving a very homogeneous result. The CV of the ten mutation positions frequency and the copy number of the tumor SNV + CNV ctDNA RM intra batch and inter-batch were less than 17% excepted positions of *EGFR*-G719S and *BRAF*-V600E. More detail of the CV value was shown in Online Resouse 4.

**Figure 4.**
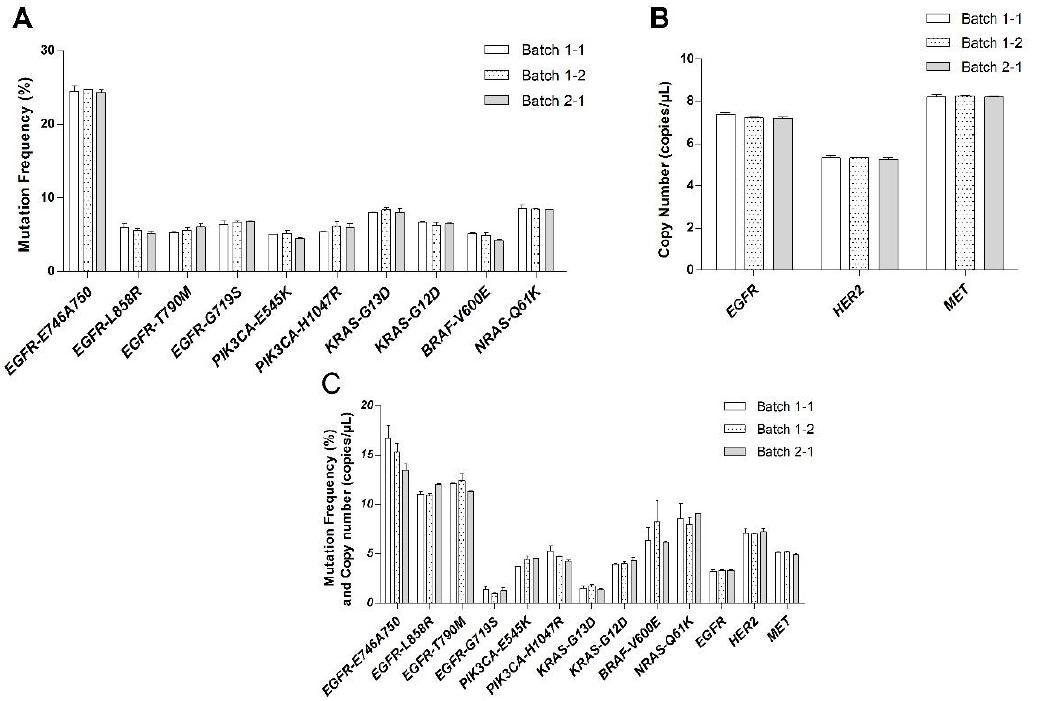
Homogenity of tumor ctDNA RMs. a. The mutation frequency of the ten mutation positions for tumor 5% SNV gDNA RMs intra batch and inter batch. Batch 1-1 and batch 1-2 are samples from same batch, while batch 1-1 and batch 2-1 are samples from different batches; b. The copy number of the three genes in tumor CNV gDNA RMs, Batch 1-1 and batch 1-2 are samples from same batch, while batch 1-1 and batch 2-1 are samples from different batches; c. The mutation frequency of ten positions and the copy number of the three genes of the tumor SNV + CNV ctDNA RMs. Batch 1-1 and batch 1-2 are samples from same batch, while batch 1-1 and batch 2-1 are samples from different batches

### Commutability of RMs

The commutability of RMs was described by the results from the four NGS panels: drug panel, pan-cancer and lung cancer-T panel as well from 1x whole genome sequencing panel (WGS). The results were shown in Figure 5. The CV value of ten mutation positions of the tumor SNV 5% gDNA RM in each panel were less than 17% except the CV value of *K3CA*-E545K (20.46%) and the thatof *KRAS*-G12D (40.27%). The CV value of the tumor CNV gDNA RM were less than 2%, showing its strong adaptation ability of different panels. However, the CV value of the tumor SNV + CNV ctDNA RM from the results of ddPCR and drug panel were not as good as its gDNA form, most of the CV value for the mutation frequency of the ten mutation positions were less than 15%, but some of them were more than 20% even up to39%. More details of the CV value can refer to Online Resouse 5.

**Figure 5.**
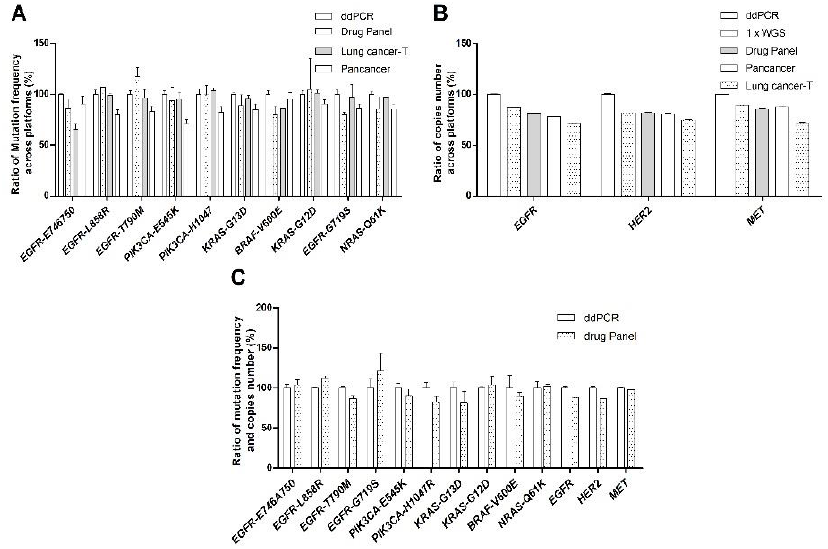
The commutity of RMs across different platform. a. Ratio of mutation frequency of ten positions from four platforms: ddPCR, drug panel, Lung cancer-T panel and pan-cancer. Data were homogeneied by the average of the mutation frequency of the ten mutationpositions from ddPCR respectively; b. Ratio of the copy number of the three genes from ddPCR, 1 x WES, drug panel, pan-cancer and lung cancer-T panel. Data were homogeneied based on the results of ddPCR respectively; c.The Ratio of the mutation frequency of ten positions and copy number of the three genes from ddPCR, and drug panel. Data were homogeneied based on the results of ddPCR respectively.

### Stability of RMs

The stability of the mutation frequency of the ten positions in the tumor 5% SNV gDNA RM and that of the copy number of the three genes in the tumor CNV gDNA RM were shown in figure 6. All CV value of the mutation frequency of the ten positions in the tumor SNV 5% gDNA RM were less than 13%, that of the copy number in the tumor CNV gDNA RM were less than 2%. These CV values indicated good stability of RMs across time. More details were shown in Online Resouse 6.

**Figure 6.**
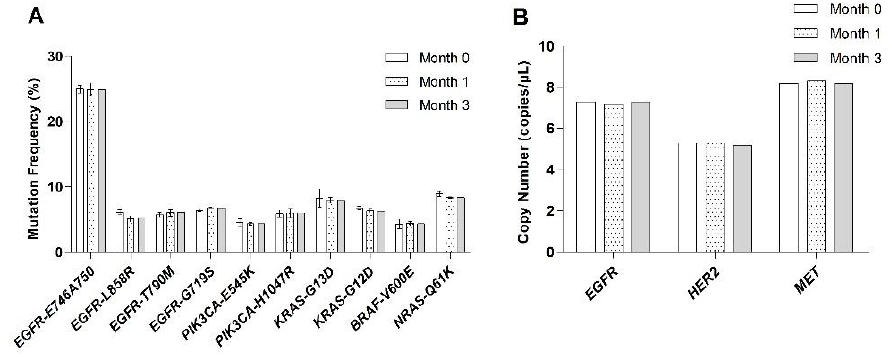
The stability of RMs across three months. a. The mutation frequency of the ten positions for tumor 5% SNVgDNA RMs. The bar chart showed the trend of the mutation frequency across three months; b. The copy number trend of the three genes among three months.

## Discussion

Development of the tumor gDNA and ctDNA RMs is significant for clinical diagnosis and tumor research. In this report, we applied droplet digital PCR to quantify the copy number of *EGFR, MET* and *HER2* genes and the mutation frequency of the ten mutation positions. The quality control of RMs was monitor based on DNA integrity and the consistence of DNA concentration. We also evaluated properties of RMs: homogeneity, commutability stability. Firstly, verified by ddPCR, the copy numbers of the three genes in tumor CNV gDNA RM remained stable at: 7.3copies/μL (*EGFR*, CV=1.5%), 5.3 copies/μL (*HER2*, CV=1.7%), 8.2 copies/μL (*MET*, CV=0.8%). The mutation frequency of the ten mutation positions in the tumor SNV 5% gDNA RM were stable for all the CV value of different batches or cross three months were less than 12%. However, the CV value of copy number and mutation frequency of SNV + CNV ctDNA RM were various. The fluctuation of the mutation frequency of different mutation positions was significant. It may due to the fragmentation of DNA, in which segment of amplification was breaks, resulting in the fluctuation. Besides, electrophoresis indicated that the DNA quality of gDNA RMs would not be affected by the mixture process according to the consistent integrity of gDNA RMs and the constant DNA concentration. Our RMs are ideal candidates for external quality evaluation and internal quality control. On the one hand, they remain stable within three months. The copy number of the three genes in the tumor CNV gDNA RM (CV value were less than 2%) and the mutation frequency of the ten mutation positions of tumor 5% SNV gDNA RM (CV value of the ten mutation positions were less than 13%) change slightly across three months. This facilitateed the monitoring of performance of detect machines or processes. On the other hand, the homogeneity of RMs makes the internal quality evaluation available. Either the homogeneity of the tumor gDNA RMs from same batch or from different batches, all of the CV valueof the copy number and the mutation frequency of the ten mutation positions were lower than 12%, except the copy number of *HER2* from second round quantification was lower than the results of first round (the fluctuation is within 25% based on ddPCR). According to their homogeneity, we can use RMs from same batch to evaluate the detection performance of different labs and compare those result. But some of the CV value of the tumor SNV + CNV ctDNA RM in different mutation positions are extremely high. They were higher than 20%. These RMs are suitable for NGS planforms of BGI, having excellent commutability, and they are good candidates for evaluate the performance of machines including ddPCR and sequencing. The copy number variation of three genes were quantified by different sequencing panels. The fluctuation of results were within ± 25% (Table 6), except the performance of CNV determination from lung cancer-T panel which did not meet the pattern of the change in panels. It was partly contributed by the unmatured bioinformatic analysis of lung cancer-T panel for CNV detection. At the same time, our RMs can reversely check the problem of BGI cancer-T sequencing panel. Besides, the CV value saw a better compatibility from ddPCR to pan-cancer sequencing panel compared with the drug sequencing panel. In drug panel, for *PK3CA*-E545K in SNV gDNA RM, the CV value of mutation frequency was 20.46, and that for KRAS-G12D peaked to 40.28. For rest of mutation positions, the CV value were less than 20%. These results indicated that our RMs were suitable for NGS planforms of BGI, with excellent commutability. Although there were some fluctuations of the mutation frequency of some mutation positions, most of mutation positions can give consistent results in different panels.

These RMs originated from clinical simples, can provides a very close genetic background and psychological background of actual cline. It can provide more accurate reference result than other RMs prepared by other methods like genetic edition.

Our RMs can be applied into a wide range of situation. On the one hand, the RMs include property of the copy number of the three genes: *EGFR, MET* and *HER2*. As a famous versatile signal transducer, *EGFR*, when it mutates or overexpresses, will induce the oncology of a number of tumors (Sigismund et al. 2018) including glioblastomas, non-small cell lung cancer(Siegelin and Borczuk 2014), head and neck, breast, colorectal, ovarian, prostatic, pancreatic cancers *etc*(Rajaram et al. 2017). Around 30% of solid tumors show *EGFR*-dependent in which tumors cannot survive and grow without the *EGFR* (Rajaram et al. 2017). Besides, *MET*, as a member of receptor tyrosine kinase (RTKs), regulates the expression of hepatocyte growth factor, plays a pivotal role in tumor initiation and progression including cancer proliferation, survival, migration, stemness, and resistance to radiation and chemotherapy(Jeon and Lee 2017; De Mello et al. 2020). This gene is found in several cancers including colorectal, breast, prostate, pancreatic cancer, and glioblastoma(Jeon and Lee 2017). Apart from *EGFR* and *MET, HER2* is a member of human epidermal growth factor receptor (Heerboth et al. 2015; De Mello et al. 2020), that strongly related with several types of cancer including carcinomas of the breast, pancreas, ovary and especially non-small-cell lung cancer (NSCLC)(Scholl et al. 2001; Vranic et al. 2021). On the other hand, the ten mutation positions involve four oncogenes: *PIK3CA, KRAS, BRAF* and *NRAS*, which enhance the range of application of this series of RMs. Mutations of these gene trigger some types of cancer including early breast cancer by *PIK3CA* (Mosele et al. 2020), colorectal cancer by *KRAS, BRAF* (Huang et al. 2018)melanomas by *NRAS* and *KRAS*(Moore et al. 2020).These genes and the ten mutation positions cover a great number types of cancer, allowing this series of RMs to be broadly applied in the tumor clinical screening and diagnosis area.

With the growing of detection methods, there will be a growing demand for RMs to have a better commutability. In our research, we mainly evaluated and compared the trains of RMs by multiple methods including ddPCR and sequencing panels of BGI. The 1x WES sequencing panel provides information of mutated genes of person to prevent the progress of malignancy. The pan-cancer sequencing panel are for broad types of cancer detection, and the lung cancer-T sequencing panel and drug sequencing panel are for monitoring the development of cancer and for evaluating the effect of treatment. Except the lung cancer-T sequencing panel, our RMs showed good stability and commutability, meaning that they can be applied into multiple purpose of cancer detection such as tumor prevention, tumor early screening and tumor monitoring. In the future, one of our works will be to construct a tumor RMs which adapts to various detection methods including ddPCR, qPCR and more sequencing panels for different purpose. Besides, our RMs cover major genes’ mutations, which can furtherly satisfy the need of cancer detection or research. But there are more mutation patterns in tumor have not been involved by this series of RMs. In the future, we would develop more RMs to facilitate the detection of mutation types like insertion, deletion, confusion and etc.

## Acknowledgements

We thanks our colleagues Qiongyao Qin and Yin Lu. They contributed this article irreplaceably. The data that support the findings of this study have been deposited into CNGB Sequence Archive (CNSA) (Guo and Chen et al. 2020) of China National GeneBank DataBase (CNGBdb) (Chen, You et al. 2020) with accession number CNP0002960.

## Notes

### Competing Interest Statement

The authors have declared no competing interest.

